# Faces of men with high serum testosterone are less attractive for women during the fertility phase of the menstrual cycle

**DOI:** 10.1101/2021.07.22.453412

**Authors:** Jaroslaw Krejza, Rafal Sledziewski, Marek Tabedzki, Rong Chen, Ewa Krzystanek, Michal Arkuszewski, Magdalena Odachowska, Kinga Kwiatkowska, Marcelina Marynowska, Maciej Rogowski, Andrzej Ustymowicz

## Abstract

The attractiveness of the human face plays an essential role in mating as it may signal the genetic suitability of a mate. The controversial ‘ovulatory shift hypothesis’ postulates that women in the fertile phase of the menstrual cycle would prefer faces of masculine men with high testosterone that signals ‘good genes’, whereas in the non-fertile phase they prefer traits signaling the willingness to provide parental care. To examine relationships between men’s testosterone and women’s preferences for men’s faces on day 13 of the menstrual cycle, 19 young women rated the attractiveness of images of the natural faces of 77 young men. Using advanced Bayesian multilevel modeling we showed that the attractiveness of men’s faces is significantly lower in men with a high concentration of serum total testosterone, even taking into account the concentration of serum estrogen in the raters. The average men’s face composited from images of 39 faces rated above pool median attractiveness rate, was slightly narrower than the average face composited from 38 less attractive faces. Our results challenge the ‘ovulatory shift hypothesis’ as faces of males with high circulating testosterone were rated as less attractive than faces of males with lower testosterone by women on the fertile phase of the cycle.

## Introduction

The attractiveness of the human face may signal the genetic suitability of a potential mate. Facial symmetry, skin’s texture, expressions of happiness, or hints of trustworthiness reflect developmental stability, health status, and ability to shape positive personal and social interactions, respectively [1–4]. Darwin suggested that individuals with genetic cues essential in the struggle for existence mate more often and produce better-adapted offspring. The favorable genotype replicated over generations leads to the development of a set of attractive phenotypic characteristics in a population of a certain geographic region. Increasing occurrences of such favorable characteristics in the population eventually develop them into dominant or alike average features over time. It shall not be a surprise therefore that an average face, created digitally by morphing many random faces together, is universally rated as attractive by the opposite sex [5].

The pace and direction of evolution of certain traits can offer a hint which features have advanced survival and reproductive successes. The remarkably rapid development of intelligence in humans signals that it has been a preferential trait. Species with larger brains, which is a proxy of intelligence, do appear to have survival and reproductive advantages [6–8]. Archeological studies of human ancestors demonstrated clearly that the volume of the skull that contains the brain expanded while the size of the face rapidly shrank [6]. Preferences for larger brains together with smaller, women-like faces taking place across cultures can explain why higher intelligence correlates with facial attractiveness, as both are highly heritable [9, 10, 11–16].

Women are more selective in choosing mates than men because they are more biologically limited in the number of offspring they can produce, and they invest more time, energy, and resources into pregnancy, nursing, and nurturing [17]. Such high biological cost is probably the main reason why women prefer long-term effective caretakers and are less receptive to sexual offers in short-term relationships [17–19]. In modern society, education, wealth, social status, and health are favored qualities as they are essentially linked to intelligence and signal the ability of a man to provide material support for their children [20, 21].

Several mating selection theories are trying to further explain women’s preferences [22]. A recent landmark hypothesis called the “ovulatory shift hypothesis” [23–28] postulates that women’s preferences for certain characteristics in men change across the menstrual cycle. Specifically, they desire men with traits signaling ‘good genes’ in the fertile phase of the cycle, whereas in the non-fertile phase they prefer ‘good fathers’ with traits signaling the presence of resources and willingness to provide parental care. Men with ‘good genes’ may not be willing to invest in a long-term relationship and may not be good providers [29, 30]. Thus, women tend to obtain these ‘good genes’ through short-term sexual affairs during the fertile phase of the menstrual cycle, and outside of a relationship with long-term effective caretakers. Such a cyclic shift of preferences supposes to justify women’s desire to obtain the best genetic quality for their children and to ensure long-term access to material resources from ‘good fathers’.

The ovulatory shift theory is being intensely disputed [31–35]. Very low rates of cuckoldry at the range from 0.73 to 3% in western societies and historical populations challenge the main concept of the hypothesis [36, 37]. The theory assumes that women can identify ‘good genes’ based on men’s phenotypic testosterone-related features such as facial masculinity and dominance [38]. The idea is based on a premise that testosterone has immunosuppressive effects and only men with ‘good genes’, which encode resistance to common pathogens, would afford to maintain a high testosterone concentration and develop masculinity features without being infected [39, 40]. Recent studies showed that dominance [41], facial masculinity [42, 43], and higher testosterone, however, are not related to superior actual health or stronger disease resistance [44, 45] and overall mating success [46]. Furthermore, there is little support for claims that health increases mating success in relatively healthy humans [47]. This negates the role of testosterone as an indicator of ‘good genes’.

Given the importance of the males’ testosterone in the mating theory, we examined relationships between men’s serum concentration of testosterone and women’s preferences for men’s faces on the fertile day of their menstrual cycle. We also tried to identify preferential face features by graphic comparison of composite images obtained from groups of attractive and unattractive men as rated by women.

## Subjects and methods

### Study design

This was a prospective analytical observational study to explore associations of the attractiveness of male faces rated by female participants in the 13^th^ day of their menstrual cycle, with male’s serum total testosterone (TT) taking into account the female’s serum sex hormones concentration. The protocol of the study was approved by the local ethical Committee. Each participant gave informed consent and was not paid.

### Population and study participants

Participants were recruited from Caucasian students of local universities. Female volunteers were included if were healthy, based on the most recent general medical examination and standard laboratory tests, did not have major health problems in the past, were not pregnant, had not used hormonal contraceptives, had regular, non-complicated menstrual cycles (from 25 to 35 days) in the last 6 months, were heterosexual, never smoked and never took drugs, had not abused alcohol and were not taking any concomitant medication. Only participants 19 to 31 years old were included to capture those with the highest reproductive potential. Male participants were included based on self-reported health status and no smoking habit. Female participants were not familiar with male participants.

We recruited 77 men (aged 20-29 years, median 22, mean±SD 22.44±1.79) and 19 women (aged 20-27 years, median 23, mean±SD 22.84±1.96).

### Hormonal status

A blood sample for measurements of 17β-estradiol (E2) and progesterone (PG) concentrations was drawn on 13th day of the menstrual cycle, after 12 hours of fasting, 72 hours abstain from vigorous physical exercises and alcohol intake. In males, a blood sample for measurements of TT concentrations was taken at the university hospital laboratory in the morning between 7 a.m. and 9 a.m. after 12 hours of fasting, 72 hours abstain from vigorous physical exercises and alcohol intake. The concentration of TT was measured according to a standard clinical practice using Chemiluminescent Microparticles Immunoassay and device by Architect ci100 (Abbott, USA), and Alinity 2^nd^ Generation Testosterone Reagent Kit (Abbott, Germany) for quantitative determination of TT. Intra-assay coefficients of variation (CV) given by the manufacturer were 3.5%, 2.5%, and 2.3% for mean TT concentration of 0.32, 2.53, and 8.43 nmol/L of in control samples, respectively. Abbott reports an inter-assay CV of 8.1%, 3.8% and 2.6%, respectively, for analogous low, medium, and high TT-level of control samples. Intra- and inter-assay CV for human serum panel was calculated at 2.8% and 3.1%, respectively, for mean TT concentration of 2.30 nmol/L. The assay is linear across the measuring interval of 0.15 to 64.57 nmol/L. We employed ARCHITECT Estradiol and Progesterone (Abbott, Ireland) kits to measure E2 and PG, respectively.

### Assessment of face attractiveness

A professional photographer took frontal view pictures of male faces in standardized studio conditions on a white background and the same lighting conditions. Clean shaved male participants were asked to maintain a neutral facial expression with their mouths closed while seated upright on a chair. They were photographed using a professional digital camera (ILCE-6500, Sony, Japan), with a horizontal and vertical picture resolution of 350×350 DPI and matrix size 6000×3376 pixels, with an exposure time of 0.01s. Flashlights were assembled to yield relatively natural lighting conditions. The distance of each face from the camera was constant at 1.8m.

We used a program, developed locally for this purpose (Inspeerity, Bialystok, Poland), to present pictures of faces on a computer screen and capture assessment of attractiveness by each female rater in an ordinal scale from 1 to 10 (10 indicates the most attractive). The raters were instructed how to operate the program and they assessed 10 phantom faces as a warm-up just before starting the trial. Thereafter, pictures of all faces were displayed in one session to each rater independently, in random order and in standardized conditions between 8:00 and 12:00am. Seven seconds were allotted for each face assessment while time flow was graphically displayed on the bottom of each display.

### Creation of composite images of faces

To combine images of individual faces and generate composite faces for comparisons, we used programs written in Python programming language (OpenCV, https://opencv.org; and Dlib, http://dlib.net). First, we detected and extracted facial landmarks (salient regions of the face, such as: eyes, eyebrows, nose, mouth, jawline – using Multi-PIE land-marking scheme from iBUG 300-W dataset from every face image [48, 49]. We applied the pre-trained face shape predictor to the face on an image, to estimate the location of 68 (x, y)-coordinates that map to facial structures on the face. The predictor was trained on the iBUG 300-W face landmark dataset. We used the facial landmark detector, which is an implementation of the One Millisecond Face Alignment with an Ensemble of Regression Trees approach [50].

In the second step, we normalized the faces and placed them in the same reference frame. Then we aligned the facial features of all input images. Subsequently, we calculated the average of all landmarks in the output image coordinates by simply averaging the x and y values. Then we used 68 feature points of each image, as well as points on the boundary of the image, to calculate a Delaunay Triangulation, which allows us to break the image into triangles. Then we aligned these regions to the average face. The vertices of the corresponding triangles were used to calculate an affine transform (the composition of transformations such as translations, scaling, and rotations). This affine transform then was used to transform all pixels inside the triangle. This procedure was repeated for every triangle in the input image [51]. To calculate the average composite image, we averaged pixel intensities of all warped images.

### Statistical analysis

We employed a statistical software (SYSTAT 12, Systat Software, USA) to perform basic data analyses. To rank facial attractiveness we used the median value of all scores assigned to the individual male face by raters, and then we classified all faces into attractive and non-attractive ones based on the median value from those assigned median values to each face. We calculated mean, median, and SD of serum hormone concentrations. We used the Shapiro-Wilk test to determine the normality of the continuous data.

We used advanced Bayesian multilevel modeling (BMM) to model the association between rate, which is the primary outcome variable (response variable), and 1) TT, and 2) TT adjusted for effects of E2, as exploratory variables. The ordinal response is assumed to originate from the categorization of a latent continuous variable. Cases (data points) from the same rater may be correlated. BMM can model the data measured on different levels at the same time, thus taking a complex dependence structure into account. In multilevel modeling, the response variable y follows a probability distribution from a family, y_i ~ D(f(*η*_i), *θ*), where y¬i is the value of y of case i, *θ* is additional model parameters. The linear predictor can be written as *η*= X*β*+ Z*μ*. In this equation, *β* and *μ* are the coefficients at the population-level and cluster-level, respectively. X and Z are the corresponding design matrices, *μ* follows a multivariate Gaussian distribution with mean zero and an unknown covariance matrix. Model parameters are (*β*, *μ*, *θ*).

In the first multilevel model, the rate was the response variable with family cumulative and TT was a predictor. We also incorporated an intercept varying by raters. That is, our model was Rate ~ TT + (1|rater). In the second model, we included TT and E2 and incorporated an intercept varying by raters.

In BMM, the inference is based on the posterior distributions which are generated by Markov Chain Monte Carlo (MCMC) [52]. Relative to non-Bayesian multilevel models, BMM can draw inference for any pattern based on posterior distributions. The classic MCMC methods converge slowly. To address this limitation, we used Hamilton Monte Carlo, which converges quickly because of its ability to avoid the random walk behavior [53]. The MCMC parameters are: 4 chains, each with 2000 iterations, and the first 1000 are warm-up.

## Results

Pooled attractiveness’ scores of male’s faces and serum TT concentrations in males are provided in table 1. Serum concentration of E2 [pg/mL] and PG [ng/mL] in female raters were within normal range at day 13^th^ of the menstrual cycle: mean±SD and range limits: 158.3±99.1, 29.0-392.0; and 0.2±0.1, 0.1-0.4, respectively.

**Table 1.** Median pooled attractiveness’ scores of natural faces of 77 males aged 20-29 years, as assessed by 19 female raters aged 20-27 years, along with males’ serum total testosterone (TT) concentrations (nmol/L).

| Parameter |  | All males | (A) More attractive<br>>median | (B) Less attractive<br><median | Difference<br>A – B |
| --- | --- | --- | --- | --- | --- |
| Score of face attractiveness* | mean | 3.99 | 4.85 | 2.92 | 1.93 |
|  | median | 4 | 4 | 3 | 1 |
|  | N | **77** | **39** | **38** |  |
| Serum TT concentration | median | 22.77 | 22.55 | 23.93 | -1.38 |
|  | mean±SD | 24.26±9.07 | 22.74±6.74 | 25.83±10.84 | -3.09 |

\*Ordinal scale from 1 to 10 (10 indicates the most attractive).

Histograms of rate, TT, and E2 concentrations are shown on figure 1. Rate and TT concentrations showed skewed distributions, thus we applied a log_10_ transformation to fulfill the condition of Gaussian distribution in modeling.

**Figure 1.**
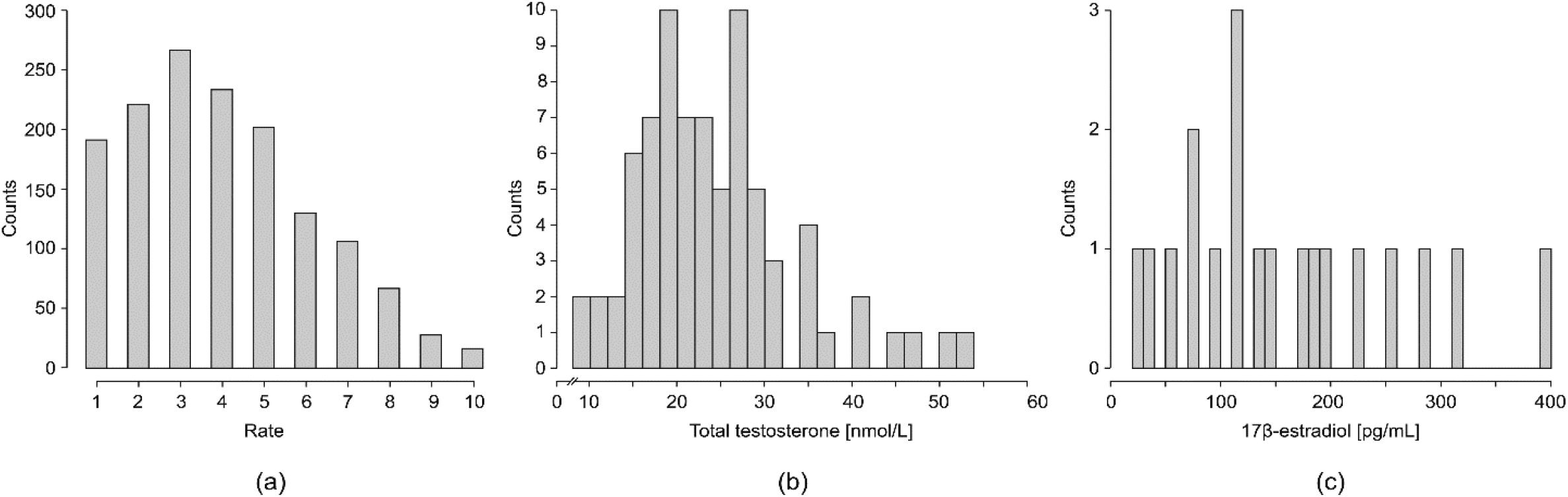
Histograms show distributions of **(a)** rate of attractiveness of 77 men as assessed by 19 women, **(b)** serum total testosterone concentration in 77 men, and **(c)** serum 17-β estradiol concentrations in 19 women

We found a significant association of males’ serum TT concentrations with attractiveness ratings. The coefficient of TT was −1.05 (estimate error was 0.3, 95% confidence interval (CI) was [−1.64, −0.48]). The higher the plasma TT concentration in male participants, the lower attractiveness score they received from female raters (figure 2).

**Figure 2.**
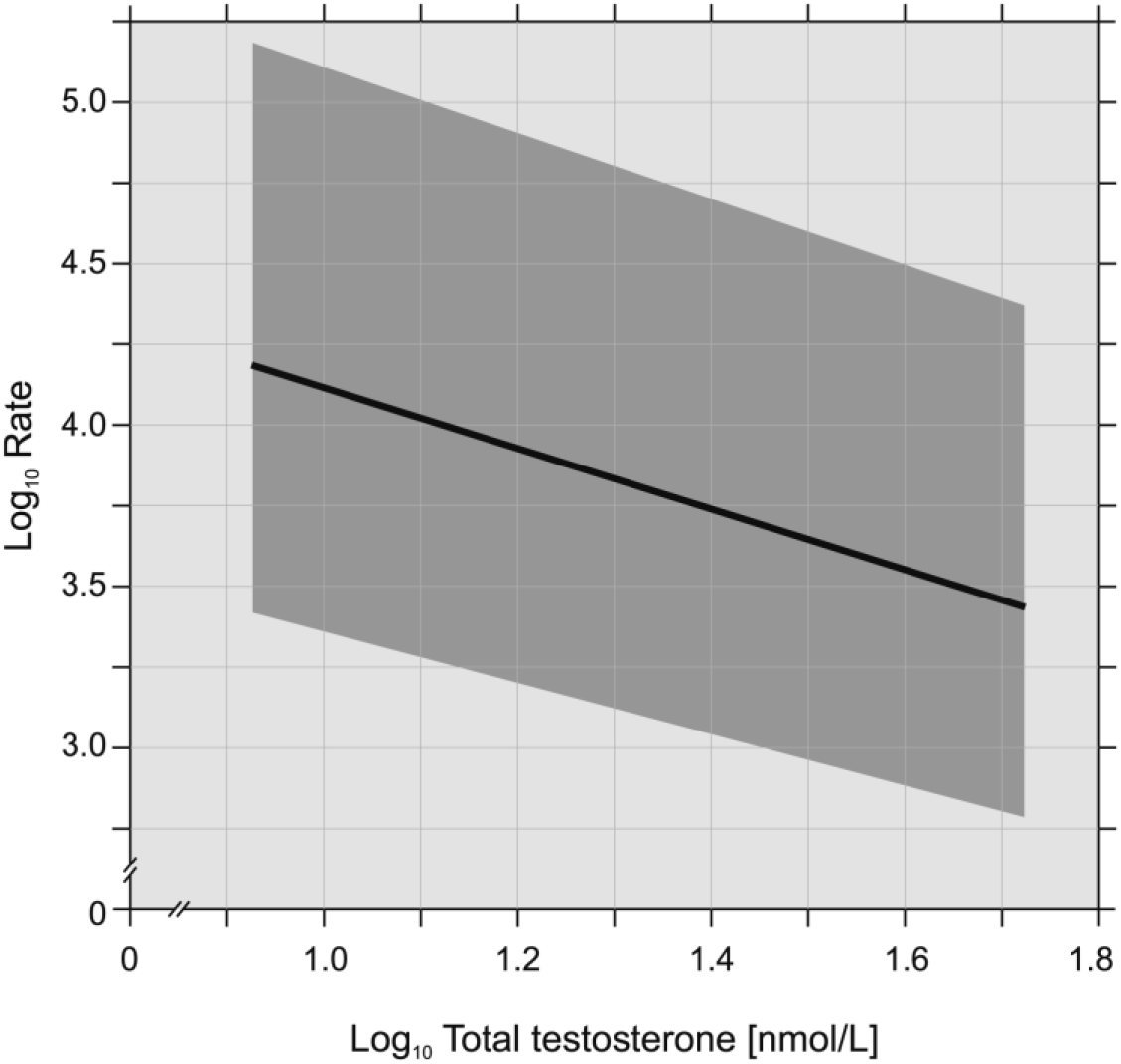
The graph shows the inverse relationship between logarithmic values of serum total testosterone concentration in 77 men and the rate of their attractiveness as assessed by 19 women.

After adjusting for E2, the association of TT with scoring was still significant. For TT, coefficient was −1.05 (estimate error 0.29, 95% CI: −1.63, −0.50). The effect of E2 was marginal, coefficient was 0.56 (estimate error 0.52, 95% CI: −0.50, 1.54). Lower scores of face attractiveness were assigned to images of faces of male participants with high TT concentration, regardless of the E2 concentration in female raters.

Three average face images composited from a) images of all 77 male faces, b) composited from 39 faces rated above pool median value, and c) composited from 38 faces rated below pool median value are shown on figure 3.

**Figure 3.**
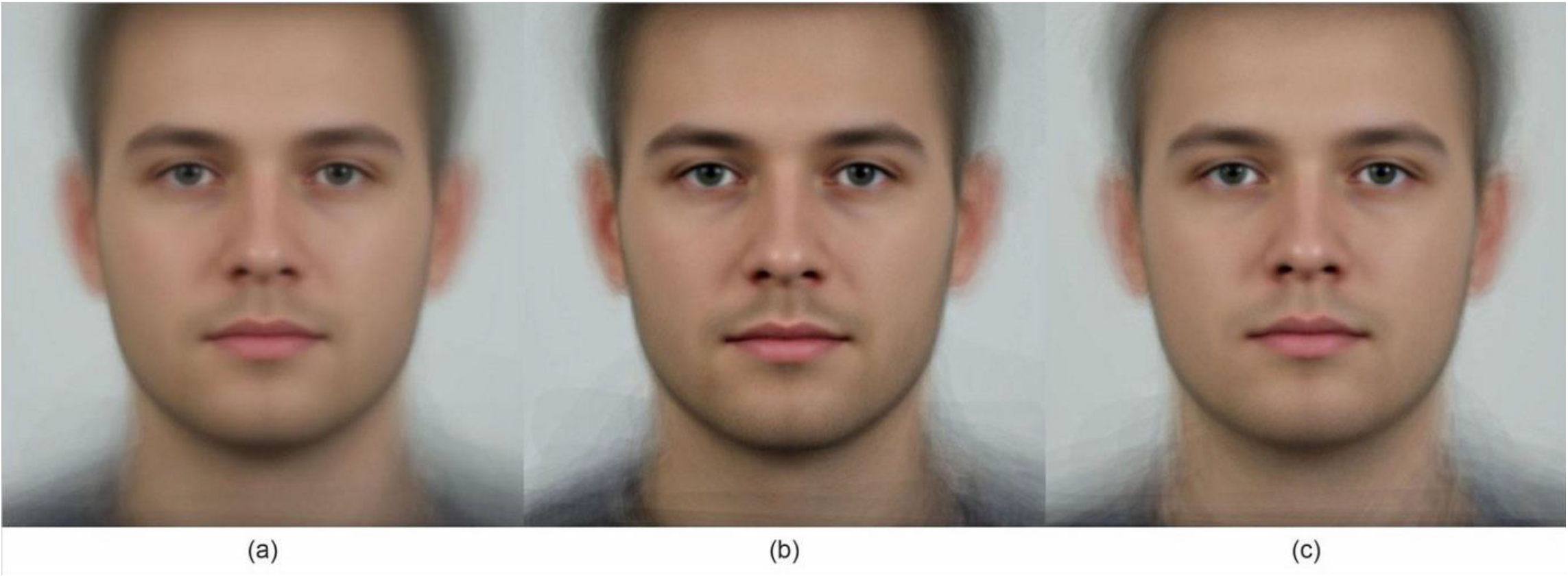
The images of a composite face of all 77 men **(a)**, of 39 men scored as more attractive than median **(b)**, and of 38 men scored as less attractive than median **(c)** by 19 female raters. Note that compared to the attractive one the less attractive composite face is slightly bigger and was scored on average by 1.93 points lower. The men from the attractive group had lower testosterone concentration than from the less attractive group.

To facilitate geometric comparison of more attractive and less attractive faces we measured the width and length of composite faces in relation to the average composite face from 77 faces (figure 4). The more attractive faces were slightly smaller than average composite faces while the less attractive were slightly bigger.

**Figure 4.**
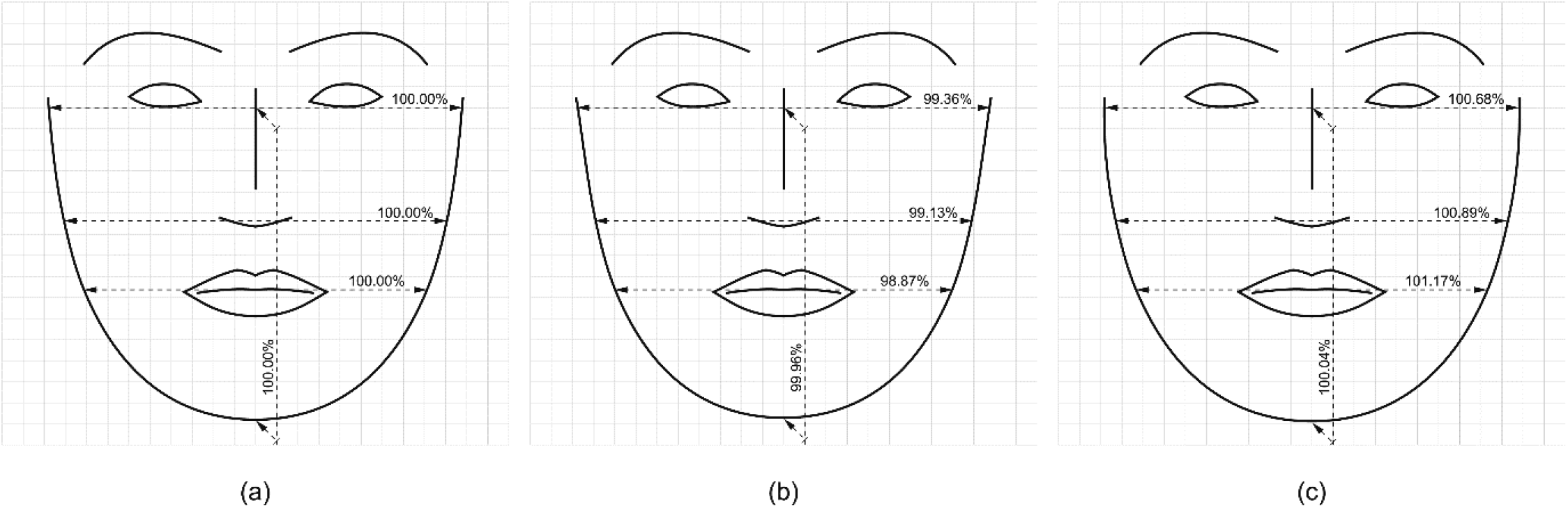
Drawings of extracted facial landmarks such as eyes, eyebrows, nose, mouth, and jawline from a composite image of faces of all 77 men **(a)**, of 39 men scored as more attractive than median **(b)**, and of 38 men scored as less attractive than median **(c)** by 19 female raters. Note that compared to the attractive one the less attractive composite face is wider by 1.32% at the level of eyes, 1.76% at the level of the lower ridge of the nose, and 2.30% at the level of mouth, and shorter by about 0.08% in midline. The average composite image of 77 men has all the measures set at 100%.

## Discussion

We have shown that images of faces of males with high concentration of circulating testosterone were rated as significantly less attractive by young women, on 13^th^ fertile day of their menstrual cycle. This result contradicts the hypothesis that women in the fertile phase of the cycle would prefer masculine, high testosterone men [54]. If such men would have been preferred by hundreds of generations of ancestral women during fertility phase of the cycle, as the theory of cyclic shift of women’s preferences suggests, androgen associated traits such massive jaw, cheekbones, and prominent brow ridges, facial and body hair, coarsen skin, low voice pitch, increase musculature, and dominant aggressive behavior would have been amplified by forces of natural intra-sexual selection [54, 55]. In contrast, several archeological and anthropometric studies provided compelling evidence that the *Homo genus*, which includes Humans, Neanderthals, and other ancestors, has experienced an evolution toward smaller faces over time, with *Homo sapiens* showing the greatest reduction in size [56, 57]. Fast shrinking face, the rapid evolution of mandibular shape and size, and rampant brain growth are suggested from analysis of numerous human fossils [57–60]. Hence the forces of evolution worked against face masculinization and favored the development of cognitive abilities.

We showed that the composite attractive male face is slightly smaller than the less attractive one. The brain can detect subtle geometric differences in faces via encoding facial features along specific axes in face space, hence facial attractiveness can be identified very quickly and reliably [61–62]. Smaller faces and lower serum TT concentration in the attractive group are mutually consistent findings, which can be related to diminished androgenic effects in earlier periods of life [63].

A broader framework of the evolution of advanced intelligence in humans connects drastic reduction of the testosterone-related aggression with diminished sexual dimorphism, which may have begun as early as 3 million years ago [6]. Aggressive behavior from childhood to adulthood can be even predicted based on testosterone-related structural brain phenotype [64]. Thus, lower testosterone favored increased tolerance between individuals for better cooperation and group survival and most likely led to social system transition from a more chimpanzee-like mating system with high levels of male–male physical competitive violence, requiring a large muscle body mass, and sexual coercion of females, toward monogamy and cooperation [65–67]. Accelerated natural selection against aggression in the last 200,000 years coincided with the further feminization of human faces as evidenced by a reduction in brow ridge projections and a shortening of the upper facial region most likely due to reduction in androgen activity on craniofacial structures [63, 67, 68]. Given that testosterone is linked to dominance, egoistic choices, aggressive and unfaithful behavior in men, the feminization of male faces may indicate preferential selection for increased social tolerance that allowed humans to work more productively [66, 69, 70]. Thus, declining androgen effects most likely “domesticated” and shrunk the human face and body size.

To the best of our knowledge, we provide for the first time empirical support for the inverse relationship between serum TT in men and attractiveness of their natural faces as scored on images by young women on their most fertile day of the menstrual cycle. Our results are in disparity with results of a study by Roney et al. who found that women prefer natural faces of men with high saliva testosterone during the fertile phase of the menstrual cycle [71]. Methodology of our study however, differs in key issues. Our samples were uniform groups of Caucasian men and women while in Roney et al.’s study the samples of men (N=37) and women (N=75) were composed of different ethnic groups. Their raters scored men’s faces in random days of their menstrual cycle, and their fertility window was merely guessed based on the calendar method and estimates of hormonal concentration from data published in other reports. Despite each woman rated only once Roney et al. did not provide crucial information on the distribution of ratings across the cycle. Especially, they did not provide how many women were in the fertile days of the cycle.

We measured concentration of TT using a standardized well-validated clinical methodology in the morning between 7 a.m. and 9 a.m., because in young men TT at 8:00 a.m. may be higher even up to 30-35% than measured in the late afternoon [72], whereas Roney et al. measured saliva ‘free’ testosterone at various times of the day. Roney et al. analyzed the scores of the physical attractiveness of faces on an ordinal 1–7 scale and correlated with testosterone concentrations, but how the scores were pooled is not given, as single woman’s attractiveness ratings of faces were regressed onto the transformed testosterone concentrations of male participants. It is puzzling, however, how pooled regression coefficients and variance from all 75 individual linear regression analyses were calculated. A simple linear regression may not be the right tool to explore the association of ordinal correlated scores of unknown distribution with testosterone values. A positive correlation among raters’ attractiveness score of the presented facial images [73], could be the result of sequentially dependent attractiveness perception or sequentially dependent response bias [74]. For example, if a rater’s scoring criterion gradually changes over time (e.g., the rater tends to give higher ratings at the beginning of the experiment and lower ratings at the end of the experiment), then the autocorrelation of the rater’s scoring criteria will lead to a positive correlation between current and previous ratings.

Peters at el. also investigated an association between saliva testosterone and attractiveness and masculinity of face and body on photographs of 119 young men [75], which were rated by two independent groups of 12 females. However, of these women, 88% were taking hormonal contraceptives that minimized any potential menstrual cycle effects on ratings. They found that testosterone was not correlated with either attractiveness or masculinity, however they implied that the correlation between testosterone and attractiveness was more likely to be negative. A small sample size of female raters, averaging scores of unknown distribution across raters, and use of Pearson coefficient to quantify associations, however, could have biased their analyses. An earlier study by Neave et al., who used natural photographs of 48 men and 36 female raters, did not find an association between salivary testosterone and attractiveness or masculinity [76]. They did not look, however, at which day of the menstrual cycle their female raters were during scoring. They also used a simple Pearson coefficient to determine relations between saliva testosterone concentrations and ratings of attractiveness.

We measured concentration of serum TT, while in experimental research investigators mostly relay on measurements of ‘free’ testosterone in saliva [71, 75, 76]. Despite speed, easy saliva collection and avoidance of stress of vein puncture, multidisciplinary clinical experts recommend serum TT as the first-line, reliable indicator of the physiological function of gonads [77]. Serum TT has low analytical variation (precision about 4% to 10%) and close correlation with calculated bio-testosterone and free testosterone [78], whereas it is uncertain if saliva testosterone represents the so-called ‘free’ fraction, because higher than expected values of ‘free’ testosterone measures in saliva are common due to binding to albumin, proline-rich proteins and steroid hormones-binding globulin [79, 80]. Hence, there is a poor agreement of saliva testosterone with serum values, even in laboratory-controlled conditions [81]. Substantial variability of measurements of saliva testosterone, related to within-person biological variations and potential pre-analytical and analytical errors [82, 83], should be accounted for when attempting to determine biologically relevant ‘least significant difference’ thresholds. Even in highly controlled conditions the threshold for salivary testosterone is large (78% to 90%) [84–86], and certainly is larger in a less controlled setting [86]. It may imply that in many previous investigations on relations of face attractiveness with testosterone statistically significant changes in salivary measures could have been obtained in the absence of meaningful biological changes.

We allowed for each rater 7 seconds time window to assess each photograph of men’s face to capture the first impression of attractiveness judgment, which is most likely rapid, automatic, and mandatory [87]. Despite that participants could have reliably judged the attractiveness of faces presented for just 13 ms [61], we selected longer time window and random display of images for each rater to minimize the systematically biased perception of face attractiveness toward faces seen up to several second before. Xia et al. demonstrated that perceived face attractiveness was pulled by the attractiveness level of facial images encountered up to 6 s after the previous image [88]. Our raters had only one task to avoid multi-task cognitive overload, which could have affected participants’ ability to intentionally form attractiveness impressions, and automaticity of impression formation. Simultaneous assessment of masculinity and scoring man’s attractiveness for a short or long term partnership would add overlay complexity for raters and would have introduced additional bias [89].

The theory of shifting women’s preferences for facial masculinity across the menstrual cycle has been intensely promoted since its introduction [23, 25, 33]. We did not assess the masculinity of participants’ faces in our study, but the smaller size of the composite face of our attractive group indicates that women prefer rather more feminized male faces. The main foundation of the disputed theory [31–33, 35, 44] however, is a premise that there is a significant association between testosterone measures and masculinity. Androgenic influence during puberty likely shapes “high-testosterone” face, which is purported to be an honest indicator of health and male fitness because testosterone enhances sexual signals, but suppresses immune function [39]. A recent meta-analysis found little evidence that testosterone suppressed immune function [42], whereas another meta-analysis identified an opposite effect - a strong suppressive effect of experimental immune activation on testosterone levels [90].

Surprisingly, there are only sparse reports that substantiate associations of testosterone with masculinity [54, 89, 91], whereas a study showed no associations between male testosterone concentrations and facial gender score [63]. The association remains an issue because of unresolved discrepancies between structural and masculinity ratings and methodological shortcomings of studies that used abstracted computer manipulated images of ‘high and low testosterone faces’ or saliva testosterone measurements [25, 75, 89, 91, 92]. Recent findings do not support preferences for male masculinity traits either at low- or high-conception probability groups of women [34, 35, 44, 93, 94]. Pound et al. [54] suggest that raters may attribute ‘masculine’ ratings to faces they find attractive irrespective of the objective sexual dimorphism, due to stereotypical associations between the term ‘masculinity’ and attractiveness. Furthermore, perceived masculinity may not correlate with attractiveness [46, 95], especially in view of an association of perception of sexual unfaithfulness with face masculinity [4].

Females’ preferences for types of male faces could differ among populations due to ethnic, socio-cultural and human development factors [9, 96]. Recent studies of samples of Slavic populations, like ours, showed women’s preferences for more feminized faces of men [9, 10, 34, 97]. Kocnar et al. reported that none of the European and non-European cultures exhibited a preference for masculinized male faces whereas feminized male faces were preferred by women in most European populations [10]. Especially, preferences for more feminized faces in men were reported in countries with high human development index [9, 10]. Perrett et al. [96] reported preferences for more face feminization of Japanese and Scottish participants compared to Caucasian North Americans faces among the raters, whereas Harris et al. have found the opposite pattern [35]. Studies comparing preferences among populations provide contradictive results, hence, the population depended differences in women’s preferences for masculinity or femininity remain an open topic.

Selection of participants of different age to conduct a research study on physical attraction to opposite sex could have an impact on the ratings and comparability of results [98]. Two studies of large cross-cultural samples found that males prefer females considerably younger them themselves and females prefer males considerable older than themselves [99, 100]. The majority of hitherto published studies are based on a sample of college male students 18-20 years old, who may still be developing their ultimate secondary sexual characteristics and face masculinity. In a study when the mean age of male participants was 18, female raters favored faces of men of higher testosterone levels [71, 101]. Male teenagers are less likely to be rated as attractive masculine men by women in their late twenties or older [96]. This could be a factor in studies, where the mean age of female raters was substantially higher than the mean age of men whose computer modified or even natural faces were used as stimuli [25, 26]. Likewise, female teenagers may give very different rates when assessing masculinity in teenagers male’s faces compared to women over 30 years old [89, 101]. Individual variation in evaluations of trustworthiness, dominance, and attractiveness, is largely shaped by people’s personal experiences and rapidity of their sexual development [98, 102]. Thus, including teenagers participants may lead to doubts whether all participants of a study are biologically and psychologically suitable to provide generalizable assessments, especially when taking into account the age disparity in sexual relationships. In our study we opted for the age matched sample.

We used natural non-edited images of male faces to explore associations between male’s facial attractiveness and serum TT. Such an approach links realistic TT measures in men with individual rates of women. Many other studies on preferences of women across the menstrual cycle in context men’s perceived attractiveness or masculinity opted to use computer simulations or heavily edited pictures or constructs of male’s face [23, 25, 34, 91, 92, 101, 103]. Certainly, computer alterations of face images to obtain a certain degree of masculinity or femininity facilitate face ratings and reduce the cost of the study, however, at the same time the differences between composite images of faces can be unnaturally high leading to believe that certain face images are more appealing to women than raw non-edited images of male faces. Ratings of images of real faces may present more appropriate assessment of women’s preferences than ratings of unrealistic computer manipulated visuals.

Several studies that use surveys of females to determine the fertile time window reported contradictive results [25, 26, 34]. The timing of women’s fertile window is unpredictable, even if their cycles are usually regular. Wilcox at al. showed that only in about 30% of women is the fertile window entirely within the days of the menstrual cycle identified by clinical guidelines—that is, between days 10 and 17 [104]. A study, in which daily values of sex hormones in female participants across the cycle were determined, showed that ovulation occurred as late as 7 days before menses, and as early as on day 8^th^ of the cycle [105]. Thus, we were fortunate enough to target accurately the fertile window as female participants were still before ovulation on day 13 of the cycle as indicated by the low level of serum progesterone concentrations.

Our sample size of female participants was rather small due to the budget constraints of the project. We used a state of the art statistical modeling, however, to reduce the probability of erroneous inferences.

In summary, we have demonstrated that young healthy women prefer images of natural faces of young men with lower concentration of total testosterone in serum on day 13 of the menstrual cycle. The size of the preferred male face tends to be smaller.

